# Insights into the structure-driven protein interactivity of RNA molecules

**DOI:** 10.1101/436279

**Authors:** Natalia Sanchez de Groot, Alexandros Armaos, Ricardo Graña Montes, Marion Alriquet, Giulia Calloni, R. Martin Vabulas, Gian Gaetano Tartaglia

## Abstract

The combination of high-throughput sequencing and *in vivo* crosslinking approaches leads to the progressive uncovering of the complex interdependence between cellular transcriptome and proteome. Yet the molecular determinants that govern interactions in protein-RNA networks are poorly known at present. Here we used the most recent experimental data to investigate the relationship between RNA structure and protein interactions. Our results show that, independently of the particular technique, the amount of structure in RNA molecules correlates with the capacity of binding to proteins *in vitro* and *in vivo*. To validate this observation, we generated an *in vitro* network that mimics the composition of phase-separated RNA granules. We observed that RNA, when structured, competes with protein binding and can rearrange the interaction network. The simplicity of the principle bears great potential to boost the understanding and modelling of cellular processes involving RNA-protein interactions.

## INTRODUCTION

Since the central dogma was proposed in 1950, the main role attributed to RNA has been to be an intermediate between DNA and protein. Yet, more than 70% of the genome is transcribed and just a small part has been found to code for proteins ^1,2^, which indicates that a major part could have unknown biological roles – if not only garbage. During the last decade many efforts have been made to develop procedures to study RNA isoforms: sequencing has been essential for detection of RNA species ^3^ and recent developments have provided a great deal of data on polymorphisms ^4^, expression ^5^ and half-lives ^6^ of all types of RNAs, which is highly informative of cellular functions and regulation. More specifically, a number of techniques reported on biological characteristics such as cellular location ^7^ or secondary structure ^8^ and characterization of the RNA interaction network (proteins and nucleic acids) is one of the most urgent challenges ^9^. In this context, computational methods are being developed to find patterns and understand features such as the structure that a transcript adopts ^10^ or to which partners are attracted ^11^.

RNA is involved in many cellular processes such as control of gene expression, catalysis of substrates, binding of ligands, scaffold of complex assemblies ^12^ and molecular chaperoning ^13^. Transcripts ability to act as a hub of cellular networks is at the centre of an active research field and has already led to the discovery of diverse ribonucleoprotein (RNP) assemblies ^14,15^. A number of membrane-less organelles have been shown to contain specific mixtures of RNAs and RBPs that are difficult to characterize ^9^. In most cases, the RNP assemblies (e.g. P-bodies, stress granules ^16^) exchange elements with the surrounding content and adapt to the environmental condition in a very dynamic way. RNA plays a central role within these phase-separated condensates: whereas a peptide of 100 amino acids can bind one or two proteins, a chain of 100 nucleotides interacts with 5 - 20 proteins ^17^. Not surprisingly, changes in the interactions within RNP granules are associated with the development of several human diseases, from neurological disorders to cancer ^18^. Importantly, regulation of RNP contacts is controlled by molecular chaperones ^18^, such as HSP70 that is a central remodelling element able to promote assembly and disassembly of RNP complexes ^19^.

In this large spectrum of activities, RNA structure dictates the precise binding of proteins by creating spatial patterns and alternative conformations and binding sites. Known complexes in which the structure of a transcript regulates protein binding include tRNAs whose three-dimensional conformation facilitates the codon/anticodon interaction ^20^ and the rRNA scaffold that sustains the ribosome ^21^. Yet, structure is not crucial only for some specific RNAs and there are several cases of nucleotide chains that play scaffolding roles: snoRNAs, for instance, are highly structured and act as a chaperone for assembly of other transcripts ^22^. The secondary structure is particularly important for messenger RNAs (mRNAs) and defines the lifecycle ^23^, recruitment of ribosomes and response against environmental changes ^23^. Of both coding and non-coding transcripts, RNA-binding proteins (RBPs) are the major regulators ^24^ and are often classified as single-stranded RNA (ssRNA) or double-stranded RNA (dsRNA) depending on their binding preferences, although this categorization is approximate.

Here we computationally evaluated the relationship between RNA structure and ability to interact with proteins demonstrating a more general and influential impact than previously reported. We linked the secondary structure to the biological function of transcripts and investigated if RNAs of a specific type or with related roles have similar structural content. Our analysis reveals a tight relationship between properties of the transcripts and their protein partners. Based on these observations, we designed an experiment to evaluate the ability of a RNA to interfere the contact network of a protein complex. Overall, our results indicate that highly structured RNAs are able to favour formation of protein assembly and remodel contact networks like a chaperone.

## RESULTS

### Highly structured RNAs bind more and stronger

With the aim of studying how the structure influences the binding of proteins, we compared human RNAs based on their secondary structure content ^8^. In this analysis we selected the least (100 transcripts, called “LS” henceforth) and the most structured (100 transcripts, “HS”) RNAs revealed by parallel analysis of RNA structure (PARS) ^8^ (**Fig. 1a**, **Supplementary Table 1**). PARS is an experimental technique that distinguishes double- and single-stranded regions of RNA using the catalytic activity of two enzymes, RNase V1 (able to cut double-stranded nucleotides) and S1 (able to cut single-stranded nucleotides) ^8^. We calculated the interactions of LS and HS sets using *cat*RAPID ^25^, an in-house algorithm that predicts the binding propensity of RBPs using physico-chemical properties (we here used 579 classic RBPs, as defined in ^11^; see **Methods**). The interaction propensity distribution (Z-score) shows that protein contacts with HS RNAs are stronger than those with LS (**Fig. 1b** and **Supplementary Table 2**). Indeed, for 501 out of 579 RBSs tested, the HS set has larger Z-score than the LS set (**Supplementary Table 2**). Respectively 34% and 18% of the HS and LS interactions show *cat*RAPID Z-score > 0 (i.e., binding ability). Thus, our computational analysis suggest that the RNA structure content is important to interact with proteins.

**Figure 1.**
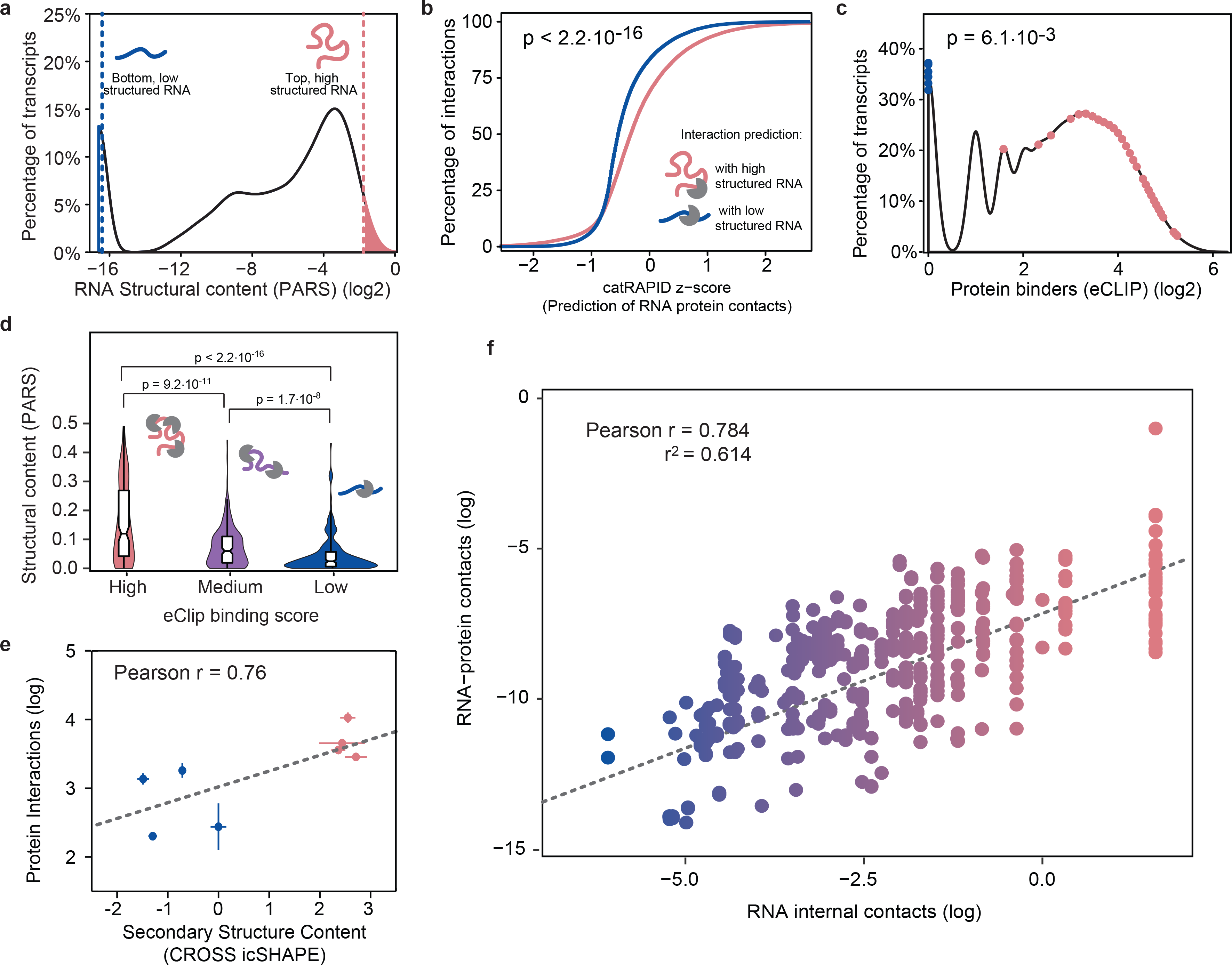
The amount of protein structure correlates with the number of interactions. **a**) Distribution of the secondary structure content of human RNA measured by PARS ^8,60^. Vertical lines indicate the top 10% cases with the lowest secondary content (LS; blue) and the bottom 10% with the highest secondary content (HS; pink). **b**) Distribution of the *cat*RAPID scores for all possible interactions with 579 canonical RBPs and transcripts with lowest and highest secondary structure content (LS and HS), respectively^11,25^. **c**) Distribution of RNAs according to their interactions with proteins as measured by eCLIP (empirical p-value of the separation is 6.1·10^−3^) ^26^. The high and low structured RNAs for (A) are mapped as pink and blue dots respectively. **d**) Violin plots showing the distribution of the PARS structural content of three groups of RBPs divided by their eCLIP binding score. High, medium and low number of RNA-protein contacts are color-coded as pink, orange and blue, respectively (see **Methods**). **e**) Correlation between structural content (CROSS predictions of icSHAPE experiments) and protein interactions of 8 transcripts revealed by protein microarrays (0.76; Pearson’s correlation). **f**) Analysis of crystals containing protein-RNA complexes reveals a trend between inter and intra-contacts of RNA chains. 196 different RNA-protein pairs analysed with different techniques and by different researchers.

To investigate whether the trend predicted by our algorithm is also observed at the experimental level, we analysed data coming from enhanced CrossLinking and ImmunoPrecipitation (eCLIP, see **Methods**), which is a technique revealing RBPs contacts on target RNAs at individual nucleotide resolution using ligation of barcoded single-stranded DNA adapters ^26^. In great agreement with our predictions, we found that the amount of double-stranded structure of each transcript correlates with the strength of protein-RNA contacts (**Fig. 1c**). It is worth to mention that the eCLIP assays favour detection of single-stranded (SS) RNA at the expense of double-stranded (DS) RNA. Importantly, the eCLIP dataset is not enriched in double-stranded RNA-binding proteins (9 out of 118 are assigned according to UniProt as DS RNA binding, 12 out of 118 as SS RNA binding, using GO annotations available ^27^), which suggests that our results are not biased by the choice of proteins used in the analysis. To further corroborate our predictions, we found that 78 out of 118 proteins interact with the HS set and 1 out of 118 with the LS (**Supplementary Table 1**). Transcripts in the LS set were found to bind between 0 and 1 binding proteins, while the HS showed a much larger span from 0 to 38 RBPs (**Fig. 1c**). We selected three sets of transcripts from the entire human transcriptome according to the protein-interacting potential as determined by eCLIP binding affinities and discovered that the propensity of RNAs to interact with proteins is proportional to the amount of RNA structure (**Fig. 1d**).

To corroborate that the observed trend is not only intrinsic to eCLIP or PARS experiments, we analysed the interactome of 8 large (>1000nt) RNAs whose protein partners have been revealed by microarray, a crosslinking-free approach ^28,29^ (see **Methods**). In parallel, we estimated the structural content of each transcript using the CROSS algorithm that was previously trained on SHAPE data ^30^ to predict the double stranded propensity at nucleotide level. Our results presented in **Fig. 1e** indicate that highly structured transcripts have more protein contacts than poorly structured transcripts, which is fully compatible with the findings presented in **Fig. 1b**.

We further corroborated our observation by an accurate analysis of the ribonucleoprotein complexes deposited in the PDB database (X-ray resolution < 2 Å; **Supplementary Table 3**; see **Methods**), which comprise 196 distinct RNA-protein pairs analysed with different techniques (X-ray, NMR) and by different researchers. Measuring the amount of RNA intra-(i.e. amount of RNA structure) and inter-contacts (i.e., amino acid) per nucleotide chain, we found a striking correlation of 0.78 between the two variables, which provides compelling evidence of their tight relation (**Fig. 1f**). Thus, independently of the experiment, the computational tool or the species we found a link between number and strength of protein interactions and RNA structural content.

### Highly structured RNAs tend to interact with proteins

The association that we describe here supports the existence of an RNA structure favouring the access to protein binding ^31,32^ (**Fig. 2a**). Literature cases supporting our observation include ribosomal RNA, for which there is a strong connection between structure and ability to scaffold protein interactions ^33,34^. Following up on this case, we wondered whether other RNA types could exploit structural regions to regulate the function of other proteins. Intriguingly, we found that the HS set is exclusively populated by protein-coding transcripts, while the LS set contains different functional classes of RNAs, such as antisense RNAs or long intergenic non-coding RNAs (lincRNAs) (**Fig. 2b**). In agreement with this observation, protein-coding RNAs are indeed the group with the largest structural content at the transcriptome level (**Fig. 2c**)^30^. Perhaps unsurprisingly, messenger RNAs and other RNA types known to interact with proteins such as snRNAs ^35^ and tRNAs ^20^ show high amount of structure, whereas RNAs targeting complementary regions in nucleic acids such as antisense and lincRNAs ^36,37^ feature the smallest amounts of structure ^38^ (**Supplementary Table 4**). Indeed, the secondary structure of mRNAs controls the translation speed ^39^ and the large difference of the coding group (**Fig. 2c**) indicates an intrinsic functional diversity.

**Figure 2.**
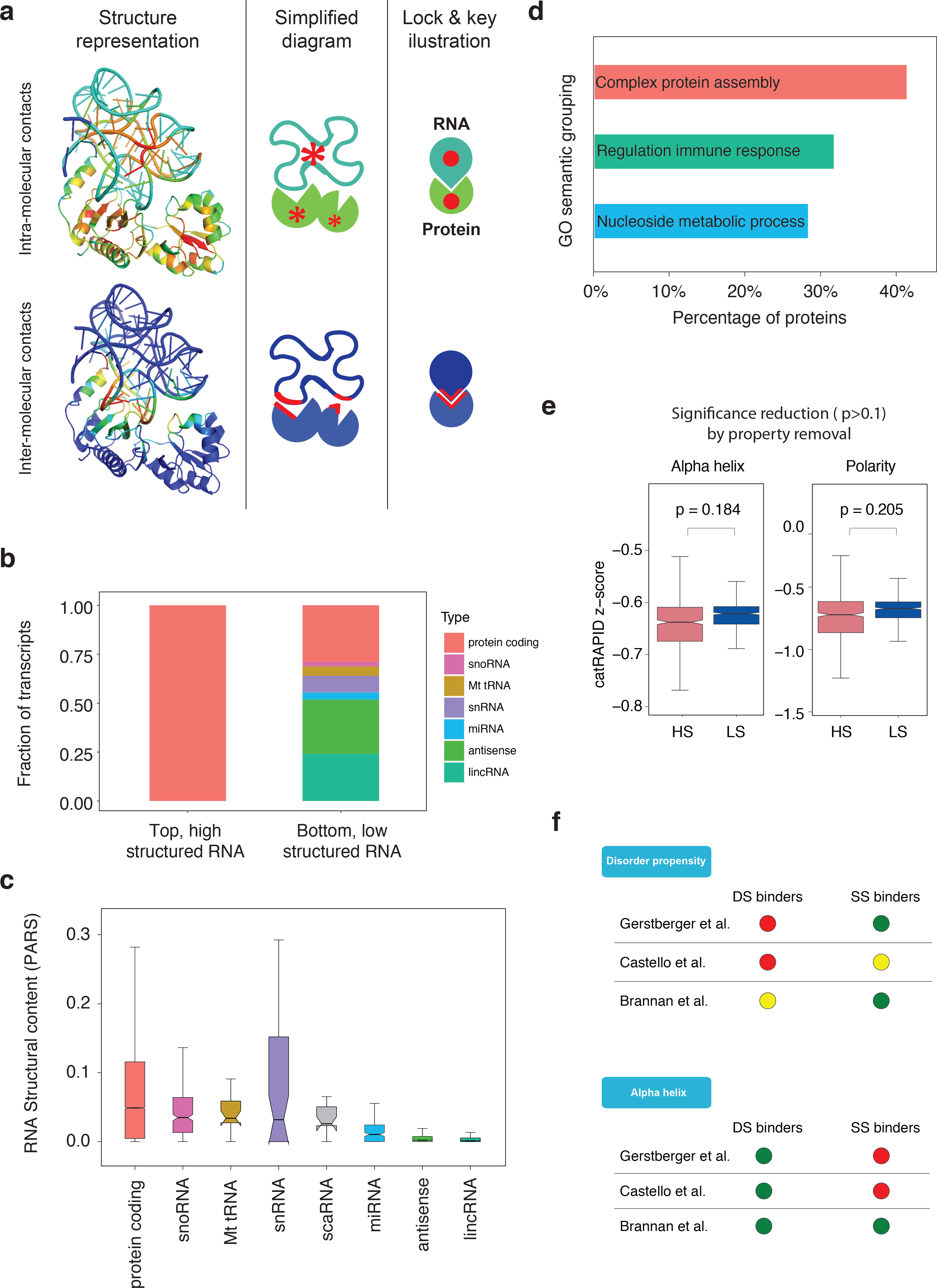
Functional footprints of the RNA structure-driven protein interactivity. **a)** Scheme showing the role of intra and intermolecular contacts in a RNA-protein complex. Top, intra-molecular contacts. Ribbon representation with red zones indicating the cores that sustain tertiary structures or local domains. Bottom, inter-molecular contacts. The simplified diagram highlights the main role: intra-molecular, sustain structures; inter-molecular, join functional elements. Contact strength, from dark blue (lowest) to red (highest). **b**) Fraction of transcripts corresponding to different RNA types according to PARS measurements ^8,60^: Left (HS) and right (LS) are the 100 transcripts with the highest and lowest secondary content, respectively. **c**) PARS structural content distribution in different RNA types (Ensembl classification). **d**) Semantic grouping of gene ontology terms associated to the HS. **e**) Changes in *cat*RAPID interaction propensities caused by removing the alpha-helix and polarity (correlating with disorder propensity, **Supplementary Table 6**) contributions abrogate the ability to distinguish between HS and LS ^11,25^. **f**) *multiclever*Machine analysis of the physico-chemical properties of three different data bases of RBPs and the set of proteins annotated in UniProt as binders of double stranded RNAs (DS) or single stranded RNAs (SS) (see **Methods**). “Disorder propensity” and “Alpha helix” are the properties showing significant difference and opposite results between DS and SS binders for at least two RBP databases (green or red indicate that DS or SS are enriched or depleted; yellow indicates no significant differences between the sets).

To further investigate the functions associated to LS and HS sets we analysed the GO terms by cleverGO ^27^. The LS set, with almost 75% of non-coding sequences, was associated with very few annotations (the current classification manly refers to coding genes) and no clustering was obtained. By contrast, several GO annotations of the HS set were retrieved and we obtained 319 terms with a Bonferroni p-value < 0.05 (see **Methods**). The analysis of semantic similarity indicates 146 terms clustered with a p-value of 0.01 (see **Methods**) (**Fig. 2d**) that can be clustered in three main groups, each covering more than a quarter of the entries: (i) Complex protein assembly (59/146), (ii) Regulation of immune response (42/146) and (iv) Nucleoside metabolic process (41/146). Intriguingly, GO terms associated with the proteins binding exclusively dsRNA (see **Methods**), are also associated with similar biological processes, such as nucleoside metabolic process and regulation of immune response (**Supplementary Fig. 1**).

Overall, our cluster analysis highlights that structured transcripts tend to interact more with polypeptides and code for proteins involved in the formation of complex contact networks, such as ‘protein complex assembly’. Accordingly, the four biological processes that involve RNA molecules interacting with proteins are: structural, hub, scaffolding and substrate. Given the relationship between RNA structure and protein interactions (**Fig. 1**), one interpretation of our results is that a high degree of control is required for genes that coordinate the activity of a large number of cellular networks ^40^. Thus, the analysis suggests a ‘recursive’ property: highly-contacted transcripts code for highly-contacting proteins, which indicates a tight level of cellular regulation ^12,41^.

### Disorder and alpha helix distinguish between double and single stranded RNA

To better understand the molecular basis for the structure-driven interactivity between proteins and RNAs, we analyzed the physico-chemical and structural parameters that allow to separate the HS and LS sets in the *cat*RAPID algorithm ^11,25^. We removed each individual parameter to estimate the impact on prediction of RNA-protein associations and found that the capacity to distinguish between protein binding to HS and LS RNAs is significantly reduced when the polarity (p=0.205) and alpha helical (p=0.184) properties were excluded (**Fig. 2e**, **Supplementary Table 5**). The property that more significantly affects the HS binding strength is polarity, which is enriched in disordered proteins ^42^ and anti-correlates with hydrophobicity (**Supplementary Table 6**) that is the most important force involved in the formation of molecular interactions ^43^. As for the alpha helical propensity, we note that helices are the most frequent structural elements involved in the formation of contacts with double-stranded regions and occur in dsRBD and Zinc fingers ^24^ (**Supplementary Table 7)**. This observation suggests co-evolution: while the RNA adopts complex shapes to expose binding regions, proteins increase their structural content. Thus, in agreement with the key lock theory, evolution have selected highly structured proteins as better interactors of double stranded RNAs ^44^.

We validated the importance of protein polarity and helical structure by comparing three datasets of well-studied RBPs ^45–47^ retrieved from UniProt as exclusively single-stranded (ssRNA, 453 proteins) or double-stranded RNA (dsRNA, 390 proteins) binders (**Supplementary Table 7**). Analysis of biophysical properties with the *clever*Machine approach ^48^ revealed that ssRNA binders and dsRNA binders are different for two properties: disorder and alpha-helix content (**Fig. 2f**). The comparison of the two sets, one against the other, indicate that RBPs binding to highly structured RNAs are structured and hydrophobic, while disordered and polar RBPs bind less structured RNAs (**Supplementary Fig. 2**). This analysis further expands what was previously reported for protein-protein networks, in which disordered regions play a central role ^40^, and identifies new rules for nucleotide base pairing with amino-acids.

### Molecular chaperones: an example of relationship between RNA structure content and protein contacts

Our analysis of the human transcriptome and across organisms indicate that highly structured RNAs are prone to interact with proteins and, in turn, code for proteins involved in biological processes with large and complex contact networks. To better investigate the structure-driven protein interactivity of RNA molecules, we focused on a class of transcripts coding for proteins interacting with a large number of partners. The natural choice for our analysis is represented by molecular chaperones, as they promote folding into the native state ^49^ and organize the assembly of ribonucleoprotein granules ^50^, thus fulfilling the ‘recursive’ property presented in **Fig. 2d**. eCLIP data ^26^ show that most of the RNAs coding for human chaperones are involved in interactions with multiple proteins (**Fig. 3a**). Confirming our hypothesis, we found a significant correlation between the protein-RNA interactions and the number of protein-protein interactions annotated in BioGRID (**Fig. 3b**). This result confirms that the transcripts bound by many RBPs code for highly-contacted proteins.

**Figure 3.**
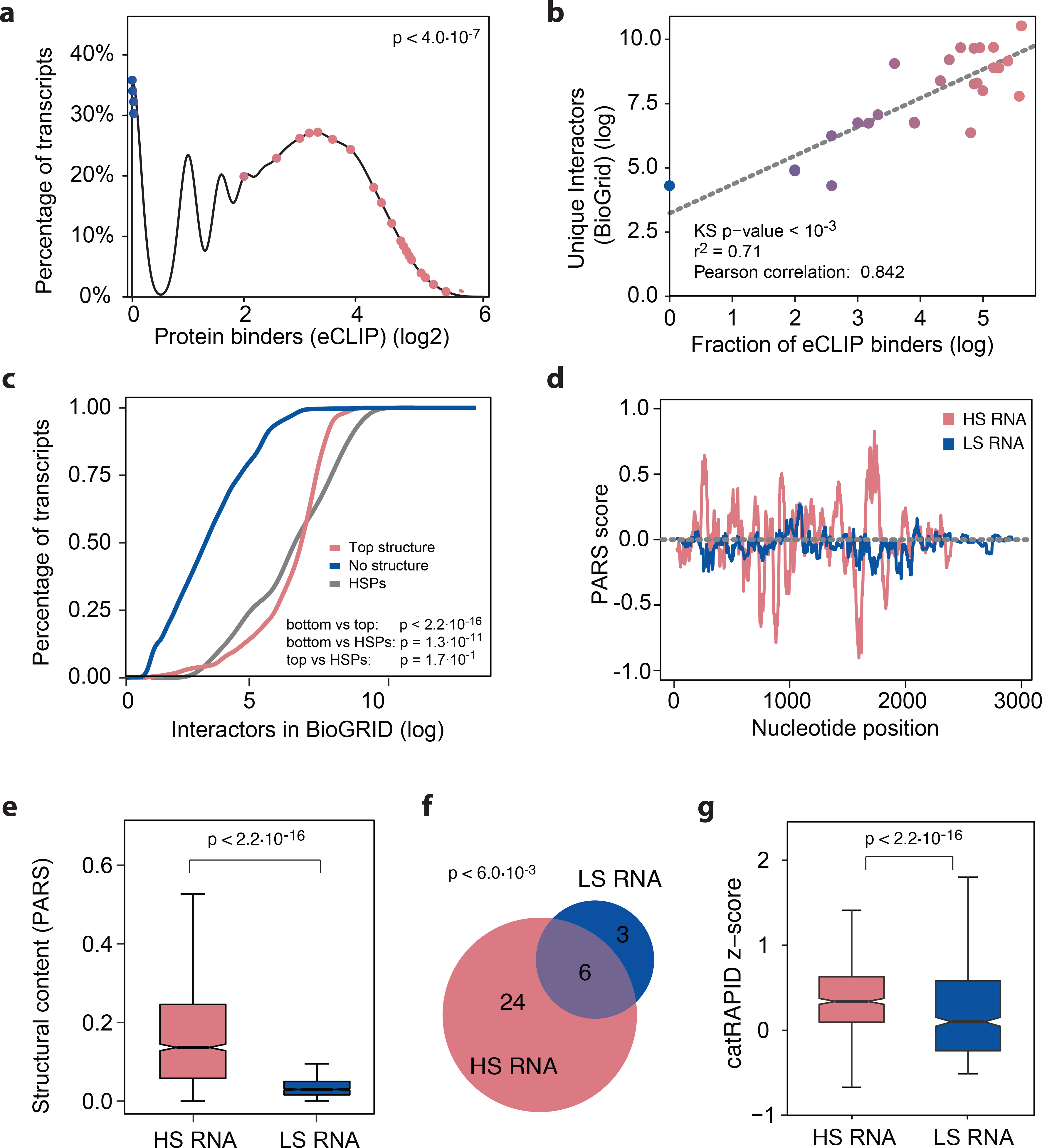
Molecular chaperones: an example of relationship between RNA structure content and protein contacts. **a**) Distribution of the fraction of proteins binding to RNAs for chaperones, as measured by eCLIP ^26^. The transcripts coding for chaperone are represented as dots blue and pink accordingly to their structural content (blue for low and pink for high, respectively). Kolmogorov Smirnov (KS0 test p-value between blue and pink dots p = 4·10^−7^. **b**) Correlation between protein contacts of RNA coding for chaperones, measured with eCLIP ^26^, and physical interactions of the corresponding proteins, collected from BioGRID. **c**) Empirical cumulative distribution function of number of physical interactors retrieved from BioGRID for three different protein sets: blue - protein products of transcripts with a PARS structural content equal to 0 ^8,60^; pink - proteins coded by transcripts with highest secondary content measured by PARS; grey - proteins corresponding to the chaperone family. blue, n=2142; red n=150; HSPs, n=31. **d**) PARS measurement of the secondary structure content of HS RNA (HSP70) and LS RNA (BRaf) transcripts. **e)** Boxplot distribution of the PARS the secondary structure content. **f**) Venn diagram showing the overlap (empirical p-value p < 6·10^−3^ computed on all the 100 eCLIP RBPs as background) between protein interactions of HSP70 and BRaf RNA. **g**) Prediction of protein binding affinity of HS RNA (HSP70) and LS RNA (BRaf) transcripts using *cat*RAPID ^11,25^.

To understand if the correlation between protein-protein and protein-RNA interactions is a general property or just associated with the chaperone family, we analysed interactions of three RNA classes: RNAs with no structure (PARS content = 0), top 100 transcripts from PARS, and RNAs coding for chaperone proteins (HSP) (**Fig. 3c**). The cumulative distribution of protein-protein contacts shows a significant difference between the “no structure” RNAs (very few interactions reported, in agreement with the results shown in **Fig. 1d**) and “top structure” RNAs (many interactions reported). The HSP transcripts have a distribution similar to the “top structure” ones. Thus, our calculations agree with the GO analyses (**Fig. 2d**) and suggest a relationship between mRNA and their coding partners: highly structured RNAs code for highly interacting proteins.

The data presented so far suggest that RNAs related either by type (e.g. miRNA, snRNA) or function (e.g. coding for chaperones) share similar structural characteristics (**Fig. 2**). Thus, we should be able to estimate differences in the interaction network of two unrelated transcripts by analysing their structural content, and *vice versa*. To test this hypothesis, we selected the transcript of HSP70: highly structured (~51% according to PARS, **Fig. 3d** and **Fig. 3e,** also with CROSS, **Supplementary Fig. 3**) and coding for a chaperone essential to regulate protein complex assemblies such as clathrin coats ^51^ and stress granules ^19,50^. As a control we chose the transcript coding for BRaf: less structured (~20% according to PARS, **Fig. 3d** and **3e**, also with CROSS, **Supplementary Fig. 3**) and encoding for an oncogene involved in transmission of chemical signals from outside the cell to the nucleus. Although there is a significant overlap between HSP70 and BRaf interactions (**Fig. 3f)**, HSP70 has more partners (30 RBPs identified by eCLIP) than Braf (9 eCLIP RBPs), which is perfectly in agreement with the structure-driven protein interactivity property.

In keeping with the trend in **Fig. 1b**, *cat*RAPID predictions show that HSP70 transcript performs stronger contacts than BRaf (**Fig. 3g**). Intriguingly, HSP70, as a highly structured RNA, codes for a protein with a higher number of interactors (244 BioGRID physical interactors) while BRaf is less structured and has a protein product binding to a smaller set of molecules (88 BioGRID physical interactors). These data suggest that an RNA with higher interactive capacity is predisposed to act as a network regulator: we can speculate that, because of its higher interactivity, HSP70 transcript could even perform a chaperone role depending on the context.

Our analysis suggests that a structured RNA, due to its higher protein-interacting potential, should affect protein interaction networks more than a less structured RNA. To validate this hypothesis, we used a chemical compound, biotinylated isoxazole (b-isox), to induce formation of a granule-like protein assembly 52,53 and incubated it with BRaf (from now LS RNA) and HSP70 (from now HS RNA) transcripts (**Fig. 4a**). We observed that the HS RNA altered the composition of the granule-like more than the LS RNA (**Fig. 4b** and **Supplementary Table 8**). A statistically significant change of concentration was determined for 29 proteins (‘released’ set) when HS RNA was added, but only for 9 with LS RNA. Clustering of the significantly changed proteins revealed that the composition in the presence of LS RNA remains similar that of the background control (‘static’ set; **Fig. 4c**). The competition of RNA with the b-isox precipitate contact network ^52,53^ could be direct or indirect (**Supplementary Fig. 4**). Yet, *cat*RAPID predictions support that this disruption is caused by a direct effect: as a decrease in the experimental stringency is associated with a drop in predictive power (**Fig. 4d**; see **Methods**). Moreover, analysis of eCLIP data indicates a higher number of contacts for the released proteins than for the static one. In agreement with our theoretical analysis the HS RNA-released proteins turned out to be significantly more polar (**Fig. 4e**).

**Figure 4.**
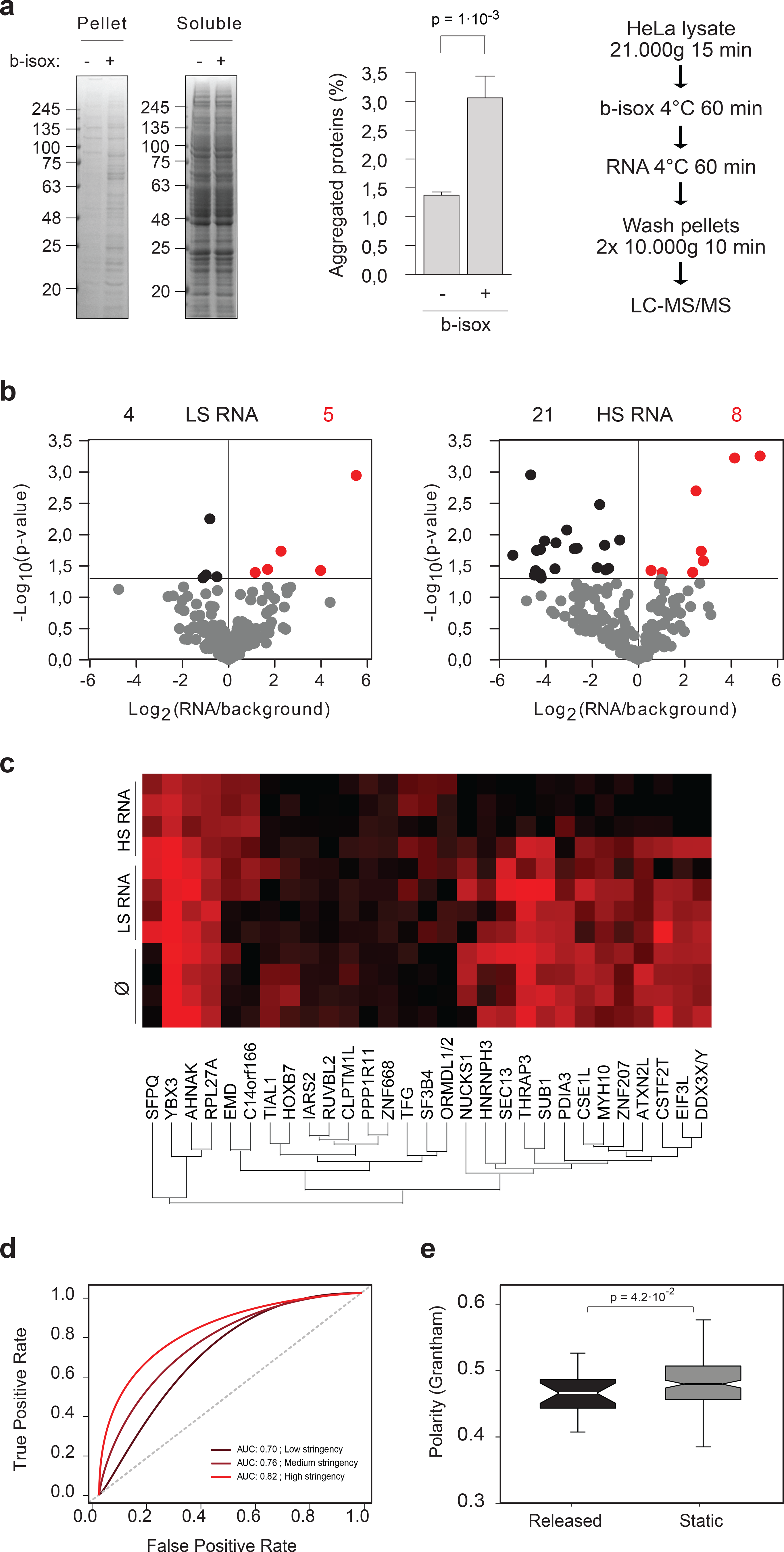
Structured RNA competes with the amyloid-like scaffold for association with cellular proteins *in vitro*. **a**) B-isox-driven aggregation of HeLa protein lysate *in vitro*. Left, coomassie-stained gels, one representative experiment shown. Center, aggregated protein intensity was quantified and the difference evaluated using two tailed t-test (p = 1·10^−3^; N=4 biological replicates). Right, experimental scheme. **b**) Volcano plots indicating the p-values (Perseus measure) of the individual protein enrichments in the b-isox assembly (N=4 independent biological replicates). The statistical significance threshold is marked by a horizontal line (see also **Supplementary Table 8**). **c**) Color-coded LFQ intensities of proteins affected by the HS RNA on a scale from black (low) to red (high). Hierarchical clustering by Perseus is indicated. For comparison, the LFQ intensities of the same proteins in control and in the presence of the LS RNA are plotted as well. **d**) *cat*RAPID predictions for positive and negative protein sets from the b-isox/HS RNA sample. **e**) Box plot of polarity distributions of proteins rescued or unaffected by the HS RNA (black or grey dots, respectively), corresponding to the right panel of **Fig. 4b** (p = 4.7·10^−2^, KS statistical test).

Overall, this experimental example demonstrates that the “recursive” trend is evolutionarily preserved and influences every level of RNA biology, from global interactome to single molecule function (**Fig. 5**).

**Figure 5.**
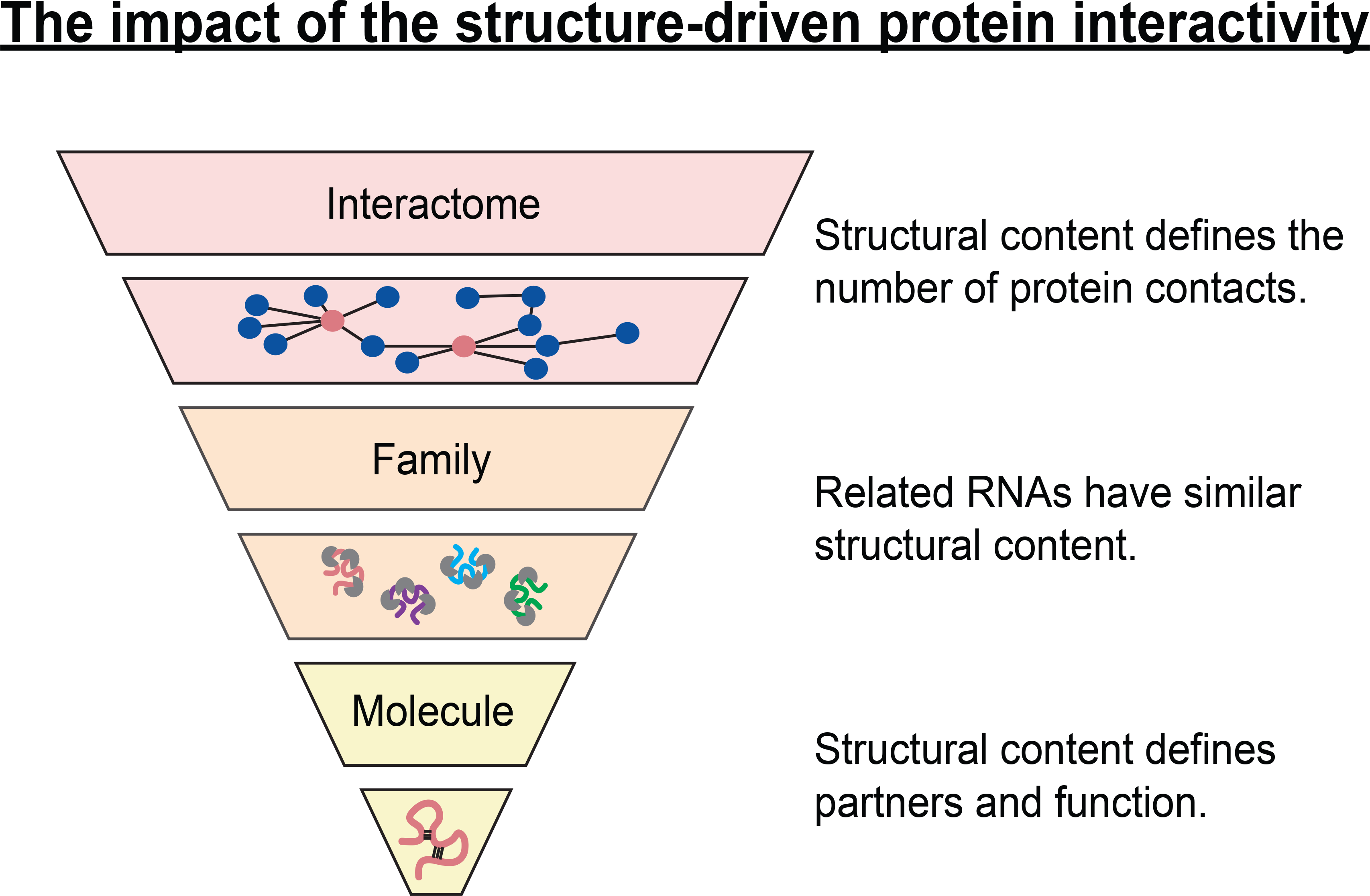
The impact of the structure-driven protein interactivity. We studied the relationship between RNA secondary structure and ability to interact with proteins demonstrating that the interaction strength correlates with the amount of RNA structure. This property, which we called the structure-driven protein interactivity, impacts every aspect of RNA life. At the interactome level, we observed that the RNA structural content defines the number of protein contacts (see **Fig. 1**). Our analysis pointed out that RNAs functionally related have similar structural content, supporting the functional impact of the structure (see **Fig. 2**). Analyzing individual RNAs we found that the structural content is linked to the number of partners as well as the function that an RNA is able to undertake (e.g. chaperoning) (see **Fig. 3** and **Fig. 4**). The structure-driven protein interactivity is an intrinsic property associated with the RNA molecule that can be traced at any regulatory level.

## DISCUSSION

RNA is often relegated to a secondary position with respect to proteins that are the major effectors of all cellular activities. However, thanks to recent experiments, it has been possible to collect information on the majority of transcripts in the cell. These advances generate big amounts of data and are revealing new functions ^9,29^. There are many questions to be addressed at molecular level to understand the full picture of RNA roles.

Here we focused on the relationship between RNA secondary structure and ability to interact with proteins. It is widely accepted that the structure of a molecule determines all aspects of its life, from stability to function ^9^. Yet, to the best of our knowledge, we are the first to show, at a transcriptome level, that the strength of protein interaction is correlated with the amount of RNA structure. We demonstrated the solidity of this principle by the analysis of crystal structures, protein micro-arrays and eCLIP data but also designing a new experimental approach. However, this observation is not completely unexpected, since lack of RNA structure is linked to more flexible and variable conformations and, thus, a shorter residence of a specific protein to a certain region. By contrast, presence of a native fold favours the formation of stable and well defined binding site that promote functional roles and, in turn, evolutionary selection. In addition, our finding agrees with the nucleotide ‘accessibility’ hypothesis defined as the probability of a protein to find its RNA motif ^31,32^. Overall, our observation identifies an intrinsic property associated to the RNA molecule that could have been exploited through to regulate the interactions between transcripts and proteins.

For many RNAs (e.g., rRNA) the structure is functionally important and therefore subjected to evolutionary selection ^8^. The correlation shown in our work indicates that structure is particularly relevant for coding RNAs and suggest an extra layer of regulation that links RNA to the protein product ^12,41^. Something similar has been previously observed in plants ^54^ for which the transcripts with high conserved secondary structure are enriched in regulatory processes in the same way as we observed for the GO ontology of the highly structured RNAs. Indeed, we observed a link between the number of protein contacts of the transcripts and the participation of the encoded protein in a large network of interactions, revealing an important level of transcriptional regulation ^29^ for these highly connected proteins ^40^. For the chaperone family the number of contacts formed by the mRNAs correlates with the contacts formed by the coded proteins.

We used a simple experimental approach to prove the relevance of RNA structure on the interaction with proteins and the functionality associated to this property. Our experiment demonstrated that a highly structured RNA, in this case the transcript coding for HSP70, is able to transform the contact network of a macromolecular complex assembly by competing with the pre-existent interactions. The main effect observed was the release of proteins from the aggregated assembly, proteins computationally and experimentally tested to be proper interactors of the HSP70 mRNA. Our finding is in agreement with previous reports indicating that RNAs are involved in RBPs assembly ^55^. Ribosomes, for instance, are known as powerful co-factors that aid in the folding of polypeptide chains as they emerge from their channel ^13^. On the one hand, RNA molecules can be regarded as chaperones assisting the assembly of proteins. On the other hand, RNA molecules are continuously handed off from one protein to another. After splicing, RNAs are escorted to the cytoplasm by proteins and delivered to the ribosome for translation. Evidence for the passage of proteins from one chaperone to another during folding provides a conceptual precedent for a chaperone action of proteins on RNA molecules^56^. Thus, there is a mutual chaperoning effect of proteins on RNA and RNA on proteins, which is likely the result of the co-evolution between the two molecules^57^. Our data suggest an intriguing activity of the HSP70 transcript as a chaperone and a connection between RNA and protein activities. Intriguingly, while the role of HSP70 protein as a protein chaperone is well documented and there are numerous reports on its binding to hydrophobic peptide domains to prevent aggregation and facilitate protein folding ^58^, very little is known about the property of its messenger RNA.

In the future kinetic analyses tracking the RNA-protein association will be needed to further elucidate to which extent protein partners actively contribute to RNA structure formation. Our findings are reminiscent of the famous lock-and-key model in the field of enzymology ^44^: the structure of both, enzyme and its substrate, are key determinants of their association. Yet, structure contributions are not trivial in the case of ribonucleoprotein associations because the combination of different nucleotides bears an obvious specificity-determining potential. While unfolded regions promote for protein-protein assembly and disordered proteins exploit short motifs to ensure high connectivity ^59^, the reduced nucleotide alphabet and its complementarity suggest that nature favors structure to connect RNAs with proteins. The observations presented here, from transcriptome to single molecule, indicate that RNA is involved in multiple levels of regulation. The complexity and diversity of protein-RNA networks reported open the avenue for the investigation of new regulatory processes.

## ACKNOWLEDGEMENTS

We thank all members of the Tartaglia’s, Vabula’s laboratory, Dr Edoardo Milanetti for the analysis of crystal structures and Dr Danny Incarnato for insights into high-throughput approaches.

The research leading to these results has been supported by European Research Council (RIBOMYLOME_309545 to GGT and METAMETA_311522 to RMV), Spanish Ministry of Economy and Competitiveness (BFU2014-55054-P and BFU2017-86970-P) and “Fundació La Marató de TV3” (PI043296). We acknowledge support of the Spanish Ministry of Economy and Competitiveness, ‘Centro de Excelencia Severo Ochoa 2013-2017’. We acknowledge the support of the CERCA Programme / Generalitat de Catalunya. Support of Spanish Ministry for Science and Competitiveness (MINECO) to the EMBL partnership.

## Author contributions

GGT and conceived the study together with the help of NSDG and RMV, AA performed the calculations, MA and GC performed and analyzed the mass-spectrometry the experiments. NSDG, RCM, GGT and AA analyzed the data. NSDG, RMV and GGT wrote the manuscript.

## References

1 Vandivier, L. E., Anderson, S. J., Foley, S. W. & Gregory, B. D. The Conservation and Function of RNA Secondary Structure in Plants. Annu Rev Plant Biol 67, 463–488, doi:10.1146/annurev-arplant-043015-111754 (2016).

2 Kashi, K., Henderson, L., Bonetti, A. & Carninci, P. Discovery and functional analysis of lncRNAs: Methodologies to investigate an uncharacterized transcriptome. Biochim Biophys Acta 1859, 3–15, doi:10.1016/j.bbagrm.2015.10.010 (2016).

3 Okazaki, Y. et al. Analysis of the mouse transcriptome based on functional annotation of 60,770 full-length cDNAs. Nature 420, 563–573, doi:10.1038/nature01266 (2002).

4 Quinn, E. M. et al. Development of strategies for SNP detection in RNA-seq data: application to lymphoblastoid cell lines and evaluation using 1000 Genomes data. PLoS One 8, e58815, doi:10.1371/journal.pone.0058815 (2013).

5 Djebali, S. et al. Landscape of transcription in human cells. Nature 489, 101–108, doi:10.1038/nature11233 (2012).

6 Tani, H. et al. Genome-wide determination of RNA stability reveals hundreds of short-lived noncoding transcripts in mammals. Genome Res 22, 947–956, doi:10.1101/gr.130559.111 (2012).

7 Tripathi, V. et al. The nuclear-retained noncoding RNA MALAT1 regulates alternative splicing by modulating SR splicing factor phosphorylation. Mol Cell 39, 925–938, doi:10.1016/j.molcel.2010.08.011 (2010).

8 Wan, Y. et al. Landscape and variation of RNA secondary structure across the human transcriptome. Nature 505, 706–709, doi:10.1038/nature12946 (2014).

9 Marchese, D., de Groot, N. S., Lorenzo Gotor, N., Livi, C. M. & Tartaglia, G. G. Advances in the characterization of RNA-binding proteins. Wiley Interdiscip Rev RNA 7, 793–810, doi:10.1002/wrna.1378 (2016).

10 Lorenz, R., Luntzer, D., Hofacker, I. L., Stadler, P. F. & Wolfinger, M. T. SHAPE directed RNA folding. Bioinformatics 32, 145–147, doi:10.1093/bioinformatics/btv523 (2016).

11 Bellucci, M., Agostini, F., Masin, M. & Tartaglia, G. G. Predicting protein associations with long noncoding RNAs. Nat Methods 8, 444–445, doi:10.1038/nmeth.1611 (2011).

12 Ribeiro, D. M. et al. Protein complex scaffolding predicted as a prevalent function of long non-coding RNAs. Nucleic Acids Res 46, 917–928, doi:10.1093/nar/gkx1169 (2018).

13 Choi, S. I., Ryu, K. & Seong, B. L. RNA-mediated chaperone type for de novo protein folding. RNA Biol 6, 21–24 (2009).

14 Maharana, S. et al. RNA buffers the phase separation behavior of prion-like RNA binding proteins. Science, doi:10.1126/science.aar7366 (2018).

15 Franzmann, T. M. et al. Phase separation of a yeast prion protein promotes cellular fitness. Science 359, doi:10.1126/science.aao5654 (2018).

16 Van Treeck, B. et al. RNA self-assembly contributes to stress granule formation and defining the stress granule transcriptome. Proc Natl Acad Sci U S A 115, 2734–2739, doi:10.1073/pnas.1800038115 (2018).

17 Chujo, T., Yamazaki, T. & Hirose, T. Architectural RNAs (arcRNAs): A class of long noncoding RNAs that function as the scaffold of nuclear bodies. Biochim Biophys Acta 1859, 139–146, doi:10.1016/j.bbagrm.2015.05.007 (2016).

18 Alberti, S. & Carra, S. Quality control of membraneless organelles. J Mol Biol, doi:10.1016/j.jmb.2018.05.013 (2018).

19 Ganassi, M. et al. A Surveillance Function of the HSPB8-BAG3-HSP70 Chaperone Complex Ensures Stress Granule Integrity and Dynamism. Mol Cell 63, 796–810, doi:10.1016/j.molcel.2016.07.021 (2016).

20 Bhaskaran, H., Rodriguez-Hernandez, A. & Perona, J. J. Kinetics of tRNA folding monitored by aminoacylation. RNA 18, 569–580, doi:10.1261/rna.030080.111 (2012).

21 Ramakrishnan, V. The ribosome emerges from a black box. Cell 159, 979–984, doi:10.1016/j.cell.2014.10.052 (2014).

22 Lestrade, L. & Weber, M. J. snoRNA-LBME-db, a comprehensive database of human H/ACA and C/D box snoRNAs. Nucleic Acids Res 34, D158–162, doi:10.1093/nar/gkj002 (2006).

23 Goodarzi, H. et al. Systematic discovery of structural elements governing stability of mammalian messenger RNAs. Nature 485, 264–268, doi:10.1038/nature11013 (2012).

24 Lunde, B. M., Moore, C. & Varani, G. RNA-binding proteins: modular design for efficient function. Nat Rev Mol Cell Biol 8, 479–490, doi:10.1038/nrm2178 (2007).

25 Agostini, F. et al. catRAPID omics: a web server for large-scale prediction of protein-RNA interactions. Bioinformatics 29, 2928–2930, doi:10.1093/bioinformatics/btt495 (2013).

26 Van Nostrand, E. L. et al. Robust transcriptome-wide discovery of RNA-binding protein binding sites with enhanced CLIP (eCLIP). Nat Methods 13, 508–514, doi:10.1038/nmeth.3810 (2016).

27 Klus, P., Ponti, R. D., Livi, C. M. & Tartaglia, G. G. Protein aggregation, structural disorder and RNA-binding ability: a new approach for physico-chemical and gene ontology classification of multiple datasets. BMC Genomics 16, 1071, doi:10.1186/s12864-015-2280-z (2015).

28 Siprashvili, Z., Webster, D. E., Kretz, M., Johnston, D., Rinn, J. L., Chang, H. Y., & Khavari, P. A.. Identification of proteins binding coding and non-coding human RNAs using protein microarrays. BMC Genomics 13, 633 (2012).

29 Marchese, D. et al. Discovering the 3′ UTR-mediated regulation of alpha-synuclein. Nucleic Acids Res 45, 12888–12903, doi:10.1093/nar/gkx1048 (2017).

30 Delli Ponti, R., Marti, S., Armaos, A. & Tartaglia, G. G. A high-throughput approach to profile RNA structure. Nucleic Acids Res 45, e35, doi:10.1093/nar/gkw1094 (2017).

31 Li, X., Kazan, H., Lipshitz, H. D. & Morris, Q. D. Finding the target sites of RNA-binding proteins. Wiley Interdiscip Rev RNA 5, 111–130, doi:10.1002/wrna.1201 (2014).

32 Hackermuller, J., Meisner, N. C., Auer, M., Jaritz, M. & Stadler, P. F. The effect of RNA secondary structures on RNA-ligand binding and the modifier RNA mechanism: a quantitative model. Gene 345, 3–12, doi:10.1016/j.gene.2004.11.043 (2005).

33 Khatter, H., Myasnikov, A. G., Natchiar, S. K. & Klaholz, B. P. Structure of the human 80S ribosome. Nature 520, 640–645, doi:10.1038/nature14427 (2015).

34 Li, F. et al. Regulatory impact of RNA secondary structure across the Arabidopsis transcriptome. Plant Cell 24, 4346–4359, doi:10.1105/tpc.112.104232 (2012).

35 Madhani, H. D. snRNA catalysts in the spliceosome’s ancient core. Cell 155, 1213–1215, doi:10.1016/j.cell.2013.11.022 (2013).

36 Chapman, E. J. & Carrington, J. C. Specialization and evolution of endogenous small RNA pathways. Nat Rev Genet 8, 884–896, doi:10.1038/nrg2179 (2007).

37 Carthew, R. W. & Sontheimer, E. J. Origins and Mechanisms of miRNAs and siRNAs. Cell 136, 642–655, doi:10.1016/j.cell.2009.01.035 (2009).

38 Yang, J. R. & Zhang, J. Human long noncoding RNAs are substantially less folded than messenger RNAs. Mol Biol Evol 32, 970–977, doi:10.1093/molbev/msu402 (2015).

39 Faure, G., Ogurtsov, A. Y., Shabalina, S. A. & Koonin, E. V. Role of mRNA structure in the control of protein folding. Nucleic Acids Res 44, 10898–10911, doi:10.1093/nar/gkw671 (2016).

40 Gsponer, J. & Babu, M. M. Cellular strategies for regulating functional and nonfunctional protein aggregation. Cell Rep 2, 1425–1437, doi:10.1016/j.celrep.2012.09.036 (2012).

41 Zanzoni, A. et al. Principles of self-organization in biological pathways: a hypothesis on the autogenous association of alpha-synuclein. Nucleic Acids Res 41, 9987–9998, doi:10.1093/nar/gkt794 (2013).

42 Wong, E. T., Na, D. & Gsponer, J. On the importance of polar interactions for complexes containing intrinsically disordered proteins. PLoS Comput Biol 9, e1003192, doi:10.1371/journal.pcbi.1003192 (2013).

43 Tartaglia, G. G. & Vendruscolo, M. Proteome-level interplay between folding and aggregation propensities of proteins. J Mol Biol 402, 919–928, doi:10.1016/j.jmb.2010.08.013 (2010).

44 Jr., D. E. K. The Key–Lock Theory and the Induced Fit Theory. Angewandte Chemie International Edition 33, 2375–2378 (1995).

45 Castello, A. et al. Comprehensive Identification of RNA-Binding Proteins by RNA Interactome Capture. Methods Mol Biol 1358, 131–139, doi:10.1007/978-1-4939-3067-8_8 (2016).

46 Brannan, K. W. et al. SONAR Discovers RNA-Binding Proteins from Analysis of Large-Scale Protein-Protein Interactomes. Mol Cell 64, 282–293, doi:10.1016/j.molcel.2016.09.003 (2016).

47 Gerstberger, S., Hafner, M. & Tuschl, T. A census of human RNA-binding proteins. Nat Rev Genet 15, 829–845, doi:10.1038/nrg3813 (2014).

48 Klus, P. et al. The cleverSuite approach for protein characterization: predictions of structural properties, solubility, chaperone requirements and RNA-binding abilities. Bioinformatics 30, 1601–1608, doi:10.1093/bioinformatics/btu074 (2014).

49 Kim, Y. E., Hipp, M. S., Bracher, A., Hayer-Hartl, M. & Hartl, F. U. Molecular chaperone functions in protein folding and proteostasis. Annu Rev Biochem 82, 323–355, doi:10.1146/annurev-biochem-060208-092442 (2013).

50 Mateju, D. et al. An aberrant phase transition of stress granules triggered by misfolded protein and prevented by chaperone function. EMBO J 36, 1669–1687, doi:10.15252/embj.201695957 (2017).

51 Sousa, R. et al. Clathrin-coat disassembly illuminates the mechanisms of Hsp70 force generation. Nat Struct Mol Biol 23, 821–829, doi:10.1038/nsmb.3272 (2016).

52 Kato, M. et al. Cell-free formation of RNA granules: low complexity sequence domains form dynamic fibers within hydrogels. Cell 149, 753–767, doi:10.1016/j.cell.2012.04.017 (2012).

53 Han, T. W. et al. Cell-free formation of RNA granules: bound RNAs identify features and components of cellular assemblies. Cell 149, 768–779, doi:10.1016/j.cell.2012.04.016 (2012).

54 Deng, H. et al. Rice In Vivo RNA Structurome Reveals RNA Secondary Structure Conservation and Divergence in Plants. Mol Plant 11, 607–622, doi:10.1016/j.molp.2018.01.008 (2018).

55 Horowitz, S. & Bardwell, J. C. RNAs as chaperones. RNA Biol 13, 1228–1231, doi:10.1080/15476286.2016.1247147 (2016).

56 Herschlag, D. RNA chaperones and the RNA folding problem. J Biol Chem 270, 20871–20874 (1995).

57 Chao, J. A., Patskovsky, Y., Almo, S. C. & Singer, R. H. Structural basis for the coevolution of a viral RNA-protein complex. Nat Struct Mol Biol 15, 103–105, doi:10.1038/nsmb1327 (2008).

58 Tartaglia, G. G., Dobson, C. M., Hartl, F. U. & Vendruscolo, M. Physicochemical determinants of chaperone requirements. J Mol Biol 400, 579–588, doi:10.1016/j.jmb.2010.03.066 (2010).

59 Tompa, P., Davey, N. E., Gibson, T. J. & Babu, M. M. A million peptide motifs for the molecular biologist. Mol Cell 55, 161–169, doi:10.1016/j.molcel.2014.05.032 (2014).

60 Kertesz, M. et al. Genome-wide measurement of RNA secondary structure in yeast. Nature 467, 103–107, doi:10.1038/nature09322 (2010).

